# PairGP: Gaussian process modeling of longitudinal data from paired multi-condition studies

**DOI:** 10.1101/2020.08.11.245621

**Authors:** Michele Vantini, Henrik Mannerström, Sini Rautio, Helena Ahlfors, Brigitta Stockinger, Harri Lähdesmäki

## Abstract

We propose PairGP, a non-stationary Gaussian process method to compare gene expression timeseries across several conditions that can account for paired longitudinal study designs and can identify groups of conditions that have different gene expression dynamics. We demonstrate the method on both simulated data and previously unpublished RNA-seq time-series with five conditions. The results show the advantage of modeling the pairing effect to better identify groups of conditions with different dynamics. The implementations is available at https://github.com/michelevantini/PairGP

## 1 Introduction

Gene expression time-series studies have become popular as they can reveal dynamics of transcriptional processes. These studies typically use longitudinal experimental designs where repeated measurements (over time) of each cell sample are collected. A common study design involves comparisons between treatments, or conditions, and the goal is to identify groups of conditions that have different gene expression dynamics. Further, to reduce variability between conditions and to increase statistical power, biological samples in different conditions are typically matched, resulting in paired longitudinal designs. Thus, it is important to take the paired design into account in the data analysis in order to reveal the true differences between different treatments.

Standard methods for longitudinal data analysis include linear mixed effect (LME) models. A number of non-linear, non-stationary and nonparametric methods for gene expression time-series have been proposed using Gaussian processes (GP) (see Supplementary materials for related research). Recently, we have developed GP based methods to implement Bayesian non-parametrics for longitudinal studies [1, 2] that can also be applied to data from paired longitudinal designs. However, posterior sampling for such models has a higher computational cost and does not scale efficiently to genome-wide studies. We propose PairGP, a GP method for paired, multi-condition longitudinal designs. The method is tested on simulated data, longitudinal gene expression data involving two treatments, and a previously unpublished longitudinal RNA-seq data from five treatments.

## 2 Methods

Each measured gene expression time-series is modeled as a combination of three components; 1) the response model, 2) the pairing model, and 3) uncorrelated random noise fluctuations. The response model is inferred from the data, so that all treatments that produce similar responses share a common response model. The pairing model is shared by all measurements coming from the same biological replicate or batch, and models the deviation from the response model. To enforce that the pairing model does not confound the response model, the sum of all the pairing model components is constrained to zero, as explained below. The model considers each gene separately. The measured gene expression *x* is transformed as *y* = log(*x* + 1) so that it can be more accurately modeled by a normal distribution. Most gene expression experiments are “hit-and-run”, where the changes are rapid in the beginning and then slow down, thus, making it a non-stationary process. To model the non-stationarity, the user is given the choice to transform the wall-clock time 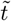 as 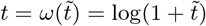. This transformation was used in all analyses reported below.

The standardized measurements of treatment (condition) *c* ∈ {1,…, *C*} and pairing *p* ∈ {1,…, *P*} is modeled as

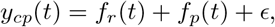

where 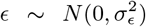. Each response effect *f_r_* is a GP with the exponentiated quadratic (EQ) kernel 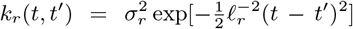, where 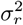 is the variance and *ℓ_r_* the length scale of response effect *r*. For each pairing *p*, the pairing effect *f_p_* is modeled with a centered EQ kernel 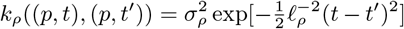, where 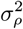 is the common variance and *ℓ_ρ_* the common length scale of the pairing effect. The centered EQ kernel has negative covariance between the pairing effects *f_p_* and *f_p′_* (*p* ≠ *p′*) to force their sum to zero, i.e, 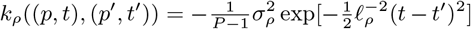 when *p* ≠ *p′* [2]. Note that the response and pairing GPs are non-stationary as the logarithmic time transformation corresponds to input-warped GPs with kernel 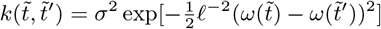. Prior distributions of hyperparameters used to analyze real data are described in Suppl. Material.

For each gene, all the partitionings of the treatments are modeled, and the one with the largest marginal likelihood (type-II) is selected as the correct response model. For example, an experiment with three treatments *c*_1_, *c*_2_ and *c*_3_ evaluates five different partitionings (models) for each gene: 1) all the three treatments have a similar response, and there is only one response model: *r*_1_ = {*c*_1_, *c*_2_, *c*_3_}; 2) treatment *c*_1_ has a different response compared to *c*_2_ and *c*_3_, and the two response models are *r*_1_ = {*c*_1_} and *r*_2_ = {*c*_2_, *c*_3_}; 3) same as (2) but with treatment *c*_2_ singled out, *r*_1_ = {*c*_2_} and *r*_2_ = {*c*_1_, *c*_3_}; 4) same as (2) but with treatment *c*_3_ singled out, *r*_1_ = {*c*_3_} and *r*_2_ = {*c*_1_, *c*_2_}}; and 5) all three treatments produce different responses, *r*_1_ = {*c*_1_}, *r*_2_ = {*c*_2_}, and *r*_3_ = {*c*_3_}.

The above method is implemented using the GPy package [3]. Instructions for the usage are available on the github page.

## 3 Results

We first tested our method on simulated time-series data with different number of treatments and a varying amount of pairing effect size (see Suppl. Material for simulation details). Comparing our model to an otherwise equal GP model but without the pairing effect shows that modeling the pairing component improves the identification of correct partitioning (Fig. 1, Suppl. Fig. 2).

**Figure 1:**
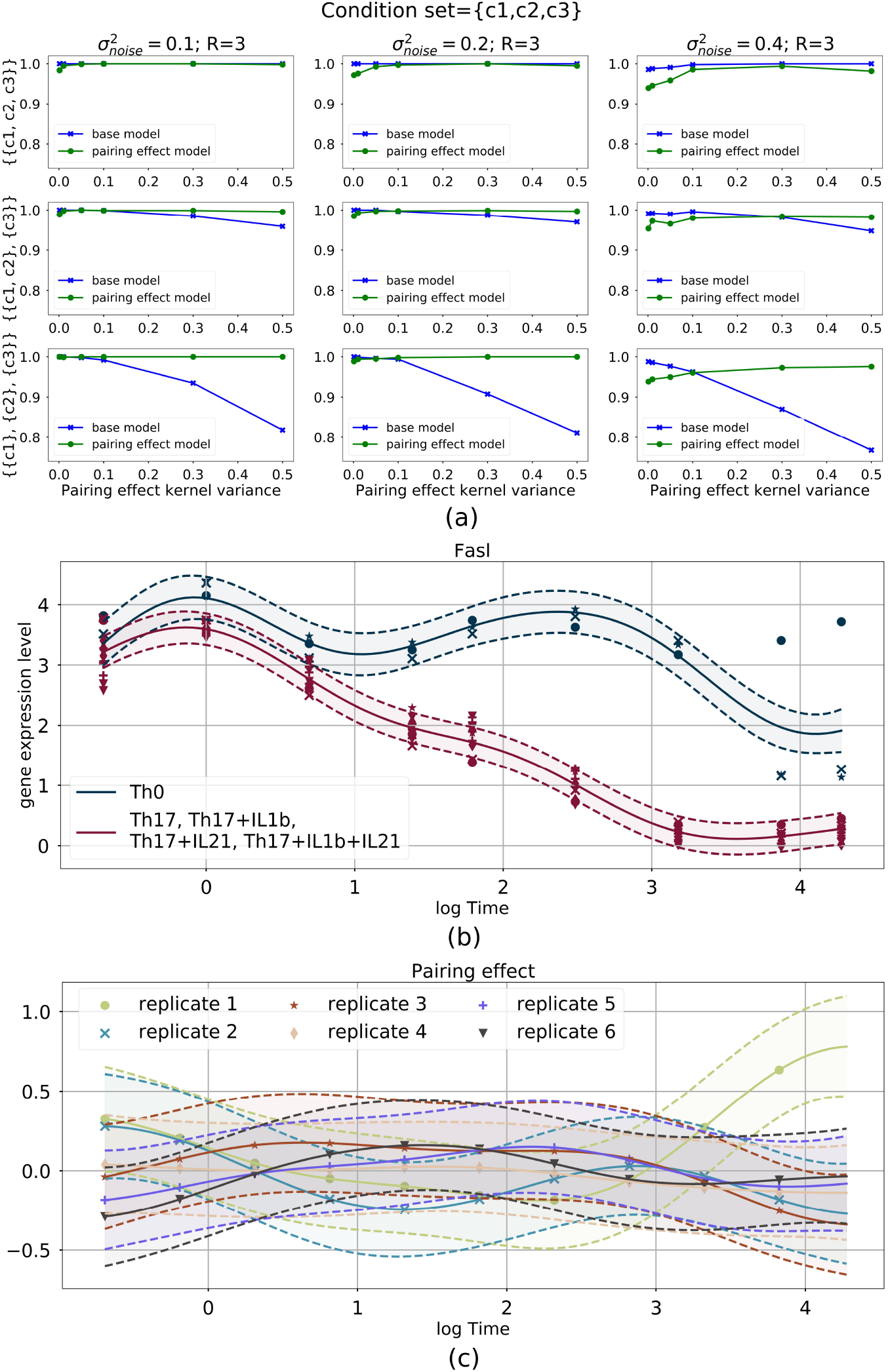
(a) Accuracy of the model inference for three treatments without pairing effect (base model) and our model with pairing effect. The accuracies (*y*-axis) are computed with simulated data where pairing effect (*x*-axis) ranges from zero (no pairing effect) to substantial (0.5). The correct partitioning of treatments for each row is indicated on left. (b) Result of the pairing effect model on the gene *Fasl*. Different colors indicate different subsets of the identified optimal partitioning and different markers represent data points coming from different replicates. (c) The pairing effect learnt from the data.

Next, we applied our method to microarray-based longitudinal gene expression data measured from activated CD4+ human T cells (Th0) and cells differentiated towards T helper 2 (Th2) cell type with three paired replicates [4]. We identified genes that respond differentially between Th0 and Th2 during the first 72 hours of differentiation (Suppl. Figs. 3 and 4, Suppl. Table 1).

We also applied our method to previously unpublished, longitudinal RNA-seq data measured from CD4+ mouse T cells that were either activated or differentiating towards Th17 lineage. Experiments include six cell cultures and five different treatments: two treatments (Th0, Th17) applied for the first three cultures and three treatments (Th17+IL1b, Th17+IL21, Th17+IL1b+IL21) for the last three cultures, resulting in two groups of three paired replicates (see Suppl. Material). Our model identifies genes that have different dynamics in different subsets of the five treatments. One example gene (Fasl) is shown in Fig. 1 and more examples are shown in Suppl. Figs. 5–6. Suppl. Table 2 summarizes how the pairing effect affects the proportion of genes detected for each partition.

## 4 Conclusions

We have implemented a GP-based model for analysis of longitudinal gene expression data that accounts for paired multi-condition study designs. Results demonstrate that our model improves the detection of correct partitioning of different conditions.

## Acknowledgements and Funding

This work has been supported by the Academy of Finland grants no. 292660 and 313271.

## Supplementary material

In this supplementary material we present the data sets that have been used in this study, details of the data simulation, some implementation details of our Gaussian process method and supplementary results.

### 1 Data

The methods have been developed generically and can be applied to any data set that has the following structure: the data set is made of *N* genes, *C* conditions (or treatments), *P* replicates and *T* time points for each gene. Thus, we have *C* × *P* time-series of length *T* for each gene. We used simulated data and two gene expression time-series data sets.

#### 1.1 Simulated data

To simulate the data, we simulated one GP for each response and one GP for each replicate pair with a fixed set of hyperparameters. To simulate the data for a specific condition *c* and for a specific replicate pair *p* we use the following formulation

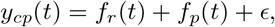

where *f_r_* and *f_p_* are GPs

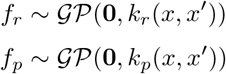

and 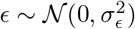 is a random noise term. Recall that the *C* treatments (or conditions) result in *R* different responses, depending on the partitioning (or model; see below), and each treatment *c* belong to one of the *R* responses. We used the same *T* = 9 time points 0.5h, 1h, 2h, 4h, 6h, 12h, 24h, 48h, 72h, as with the real data (see below). Once the kernel hyperparameters are fixed, then to simulate the data for a gene with *C* conditions and *P* replicates we simulate one realization from *f_r_* for each response effect, and one realization from *f_p_* for each replicate pair and we combine them together with additive noise as shown above. As a result, we obtain *C* × *R* × *T* time points for each simulated gene.

In our settings, we used the EQ kernels with lengthscale *l_r_* = 1.0 and variance 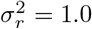 for the response kernel *k_r_*, and *l_p_* = 1.0 and 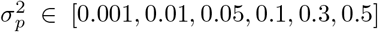 for the pairing effect kernel *k_p_*. If the individuals that are participating to the study are, for example, studied in a controlled environment, such as laboratory mice, then the variation between individuals is expected to be smaller compared to studies done with humans. Therefore, we decided to simulate data with different levels of pairing effect variance, aiming to cover several possible values. Additionally, for the random noise variance 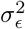 we used the set of values [0.1, 0.2, 0.4]. Similarly as for the pairing effect variance, the noise variance can also vary depending on the experiments. Thus, we want to assess the performance of the pairing effect model on simulated data as a function of the pairing effect variance and the noise variance, and compare the results to those obtained with the base model on the same data. We simulated gene expression time-series data with 3 and 4 conditions and 3 replicates with all the combinations of parameter values described above.

To evaluate the accuracy on simulated data we generated several simulated data sets, which is composed of genes generated according to each of the possible partition of the conditions. In particular, we used all the combinations of pairing effect variance, noise variance and number of conditions mentioned above to analyse how the pairing effect model behaves under several different settings. A total of 1000 genes for each partition have been simulated in each data set. However, some partitions of the conditions are *de facto* the same, for example, simulating data as {{*c*_1_}, {*c*_2_, *c*_3_}} or as {{*c*_1_, *c*_2_}, {*c*_3_}} is equivalent. Therefore, we simulate data for 3 partitions for a data set with 3 conditions, and 5 partitions for a data set with 4 conditions. Importantly, however, the model during the inference process can still choose between all the possible partitions of the condition set.

#### 1.2 Human T-helper cell differentiation data

The first data set contains gene expression time-series data from human CD4+ T cells measured using microarrays originally published in [1]. We use data from two treatments measured at time points 0.5h, 1h, 2h, 4h, 6h, 12h, 24h, 48h, 72h. Th0 condition (or treatment) corresponds to activation of naive CD4+ T cells, and Th2 corresponds to activation and differentiation of naive CD4+ cells towards T helper 2 (Th2) lineage. Both conditions (across all timepoints) are measured from three cell cultures that correspond to three biological replicates that are paired across the conditions. Microarray data is RMA preprocessed as in [1] and further standardized.

#### 1.3 Mouse T-helper cell differentiation data

The second data set has been collected from laboratory mice, and it has a total of five treatments and six cell cultures (i.e., biological replicates). The experimental details are as in [2]. Th0 treatment corresponds to activation of naive T cells. The other four treatments are Th17, Th17+IL1b, Th17+IL21, Th17+IL1b+IL21. Th17 corresponds to activation and differentiation of naive CD4+ cells towards T helper 17 (Th17) lineage. Th17+IL1b, Th17+IL21, Th17+IL1b+IL21 treatments corresponds to simultaneous activation and differentiation of naive CD4+ cells towards Th17 lineage and treatment with interleukin 1 beta (IL-1*β*), interleukin 21 (IL-21) and combination of IL-1*β* and IL-21 (with concentration 20 ng/ml) (R&D Systems), respectively. Experimental data for the treatments Th0 and Th17 have been measured from the first three replicates (cell cultures), using a paired design. Experimental data for the other three treatments have been measured from the other three replicates (cell cultures), again using a paired design. Cells are sampled for gene expression analysis at nine time points: 0.5h, 1h, 2h, 4h, 6h, 12h, 24h, 48h, 72h. Sequence reads were mapped with TopHat to mouse mm9 genome as well as to Ensembl transcriptome. After the alignment, the number of reads that mapped to each gene were summarized using HTSEQ-count tool. The raw RNA-seq data used in this manuscript will be made available upon publication via Gene Expression Omnibus (GEO).

#### 1.4 Data standardization

After quantification of gene expression count data from the RNA-seq data, the expression data is further log-transformed. Microarray and RNA-seq data are standardized before analysis.

### 2 Methods

#### 2.1 Previous methods

Gene expression microarray and RNA-seq techniques allow quantitative, genome-wide analysis of gene expression levels. A number of software tools are available for statistical analysis of gene expression data measured by microarrays (e.g. LIMMA [3]) and RNA-seq (e.g. DEseq [4] and edgeR [5]). These tools rely on linear and generalized linear models, use empirical Bayes to share information between genes, allow modeling complex experimental designs, and support testing a variety of hypothesis, but are not designed for longitudinal studies that involve repeated measurements of individuals over time. Standard methods for longitudinal data analysis include linear and generalized linear mixed effect (LME) models, as implemented in e.g. lme4 package [6]. Bayesian alternatives for modeling gene expression time-series data have been proposed e.g. in [7, 8] that also support non-Gaussian likelihood models. Methods of gene expression time-series data analysis include also *lmms* [9] and ImpulseDE2 [10]. The former is based on linear mixed models and ANOVA log likelihood ratio tests, while the latter is based on an impulse model as a continuous representation of temporal responses.

A number of non-linear, non-stationary and non-parametric methods for gene expression time series have been proposed using Gaussian processes (GP). Yuan was among the first who used GPs to model gene expression time course data [11]. A number of improved methods have been proposed, such as methods that can account for outliers [12], a method for analyzing multiple conditions [13], methods that identify time intervals of differential expression [12, 14], and methods for accounting time delays between replicates and non-Gaussian likelihood models [15]. However, none of these tools can account for paired experimental designs that are commonly used in biological studies. Similar ideas have been proposed in the context of GP-based clustering of time-series data [16], where authors propose a hierarchical GP regression model. Nonetheless, the effects in [16] are not across replicate pairs but, instead, a different replicate effect is learned for each individual condition. To that end, Spies et al. [17] provide an extensive review of a large selection of methods proposed in the literature for time course data.

Recently, we have developed GP based methods to implement Bayesian non-parametrics for longitudinal studies [18, 19] that can also be applied to data from paired longitudinal designs. However, posterior sampling for such models has high computational cost. We propose a non-stationary GP method for paired, multi-condition longitudinal designs that provides efficient analysis for genome-wide studies.

#### 2.2 Model selection

Given that an experiment contains *C* treatments, they can be partitioned into *B_C_* different partitionings (or models), where

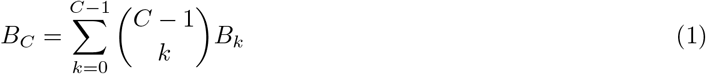

is the Bell number. For example, Bell number for 3, 4 and 5 treatments are *B*_3_ = 5, *B*_4_ = 15, and *B*_5_ = 52. For each partitioning, we evaluate the marginal likelihood

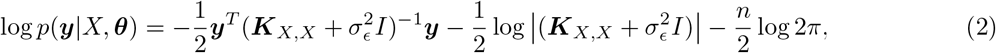

where 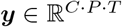 contains the standardized gene expression data for a gene from all *C* treatments, *P* replicates and *T* time points, *X* = (***x***_1_,···, ***x***_*C·P·T*_) contains the explanatory covariates (treatment *c*, replicate *p* and time point *t*) for each measurement, ***θ*** is a vector containing all the kernel hyperparameters, ***K**_X,X_* is the sum of the response covariance matrix and the pairing covariance matrix defined by the centered EQ kernels, 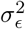 is the Gaussian random noise variance, and *n* = *CPT*. An example of the covariance matrix ***K**_X,X_* and its components ***K**_r_* and ***K**_p_* are shown in Figures 1.

**Figure 1:**
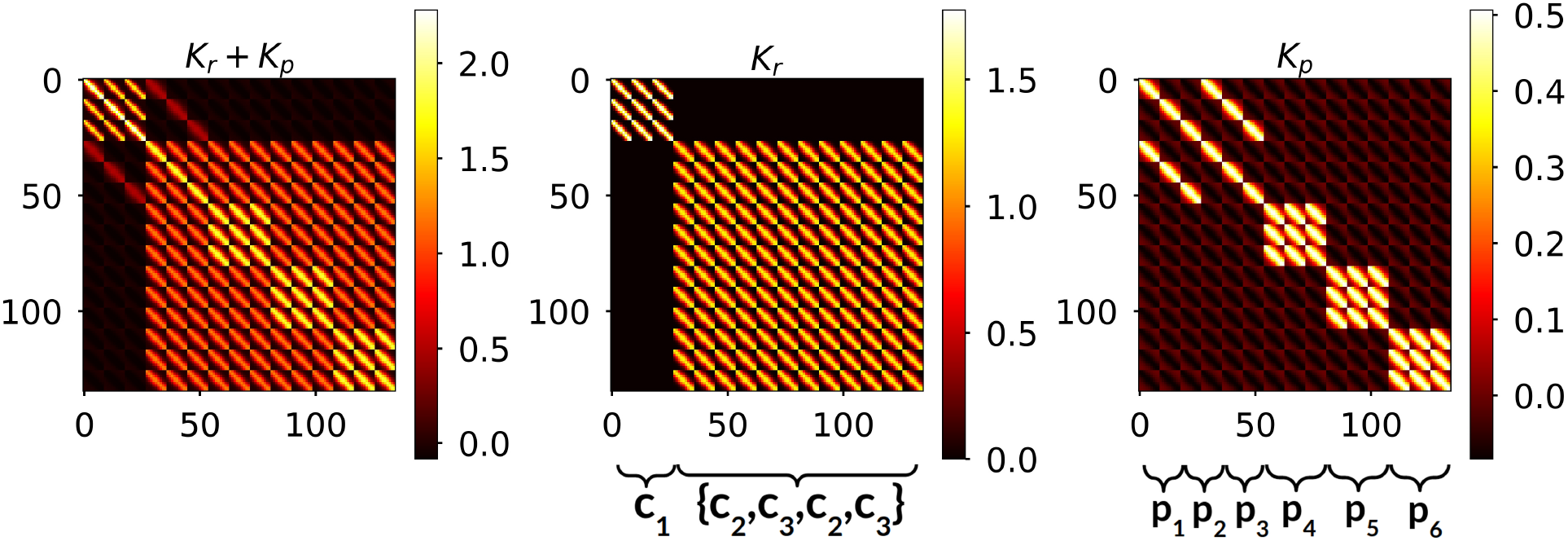
An example of the covariance matrices. (left) the combination of covariance matrices ***K**_X,X_* = ***K**_r_* + ***K**_p_* without centering. (middle) the response covariance matrix ***K**_r_*. (right) the pairing covariance matrix ***K**_p_* without centering. In this example, there are data from 5 different conditions and conditions are assumed to be partitioned into 3 response models *r*_1_ = {*c*_1_}, *r*_2_ = {*c*_2_, *c*_3_} and *r*_3_ = {*c*_4_, *c*_5_}.

We call the model presented above the pairing effect model. To assess the performance of this model, we compare it against the base model. The base model is obtained by optimizing one GP regression model for each possible subset of the condition set, and then combining the score of these models to have a score for each partitioning of the condition set. In other words, the log marginal likelihood log *p*(***y***|*X, **θ***) of the models of different subsets is summed up to obtain the score for the partitioning that corresponds to the set of considered subsets. In the base model, we use EQ kernels which model the response functions, but not the pairing effect. In the pairing effect model we standardize the data of all the conditions together, while in the base model we standardize separately the data of the sets of conditions that corresponds to the different response functions.

#### 2.3 Prior distribution for kernel hyperparameters

Typically, the hyperparameter optimization is done by maximizing the log marginal likelihood of the model. If one has prior information about the hyperparameters, then in a hierarchical structure one can also impose prior distribution on the hyperparameter, also called hyperprior. The kernel choice for the GP regression models is the exponentiated quadratic (EQ) kernel. This means that we can define hyperpriors for the variance *σ*^2^ and the lengthscale *ℓ*. The assumption that we have on the data are essentially two:

- the lengthscale parameters for the response effect kernels should be relatively high as higher lengthscale parameters imply smoother functions.
- The variance parameter for the pairing effect kernel must be relatively small; the magnitude of the pairing effect cannot be as high as the response effect, but it should just represent slight variation around condition mean that is associated with the different replicates. The only exception to this is when a gene is silent. In this case, the variation in the gene expression over time is almost 0, thus the variation due to the effect introduced by the different replicate can be potentially higher.

We use the log-Gaussian distribution log-Normal(*μ, σ*^2^) with *μ* = 0.5, *σ*^2^ = 0.5 as hyperprior distribution for the lengthscale of the condition effect, exponential distribution Exp(λ) with λ = 2 for the pairing effect variance and log-Gaussian distribution log-Normal(*μ, σ*^2^) with *μ* = 0, *σ*^2^ = 0.5 for the pairing effect lengthscale. We do not use here any hyperprior distribution on the noise variance 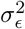.

The optimization is done w.r.t. to the following objective function

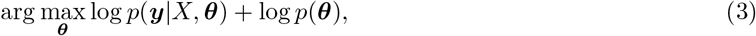

where *p*(***θ***) represents the hyperpriors. We use the above prior distributions for kernel hyperparameters when analyzing real microarray or RNA-seq data and optimize the above objective function. For simulated data we ignore the hyperpriors and optimize the standard marginal likelihood, i.e., log *p*(***y***|*X, **θ***). We use the gradient-based method L-BFGS-B [20] for the optimization. The optimizer is run for a maximum of 1000 iterations with tolerance for deciding convergence equals to 1*e*^-5^.

### 3 Results

#### 3.1 Simulated data

**Figure 2:**
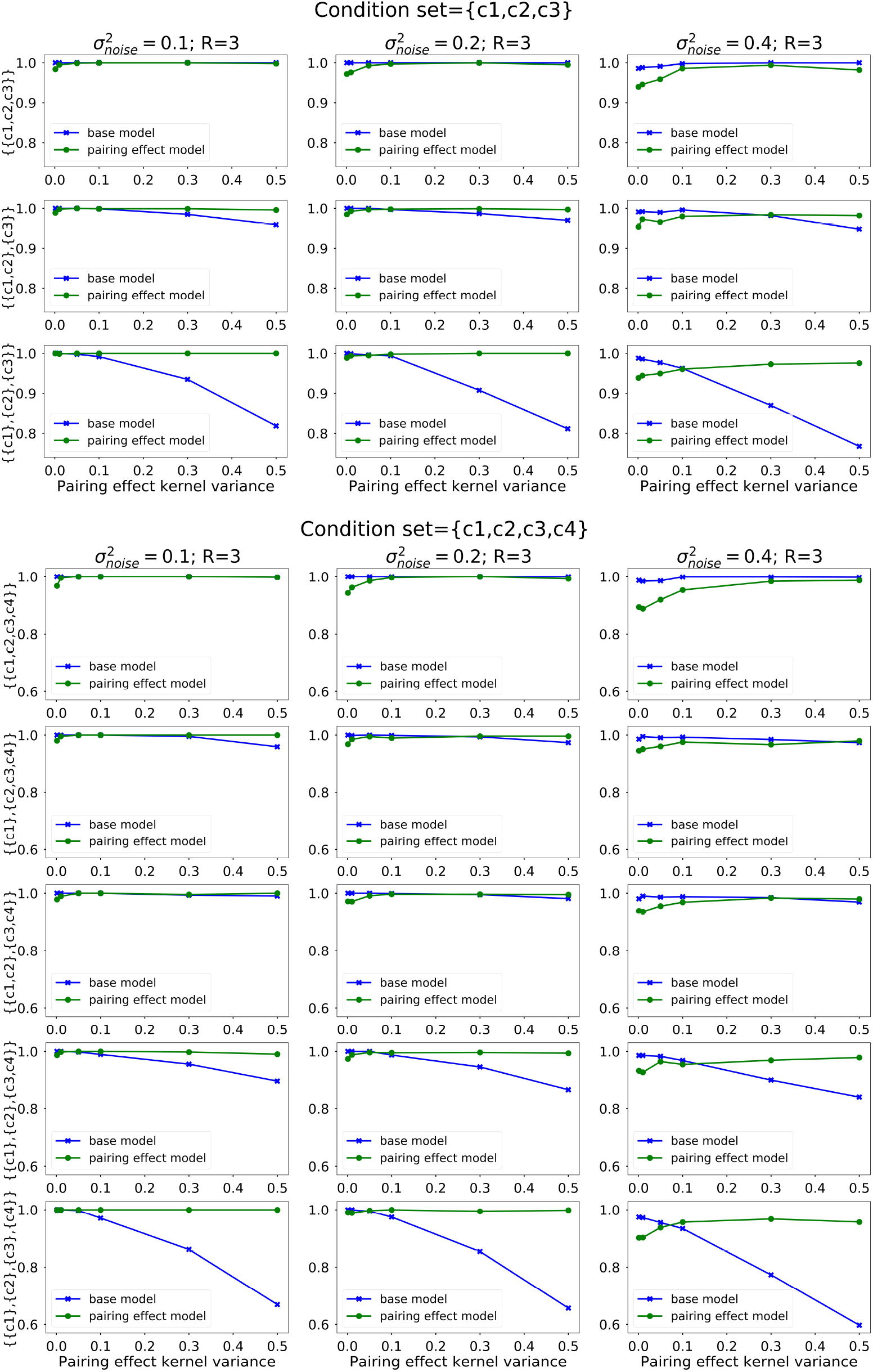
Accuracy of inferring the correct partitioning of conditions as a function of the pairing effect variance and the noise variance, obtained through simulated data using (top) 3 conditions (bottom) 4 conditions.

#### 3.2 Microarray data

**Table 1:** Proportion of partitions obtained by fitting the base model and the pairing effect model on the genes from the human T-helper cell gene expression data set. Thus, when taking into account the paired design of the experiment, 30.4% of the genes were found to be differentially expressed between Th2 and Th0 cells, whereas 24.9% of genes were differentially expressed when only the response effect was modelled.

| Partition | Base | Pairing |
| --- | --- | --- |
| $\{\{\text{Th0}, \text{Th2}\}\}$ | 75.1 | 69.6 |
| $\{\{\text{Th0}\}, \{\text{Th2}\}\}$ | 24.9 | 30.4 |

**Figure 3:**
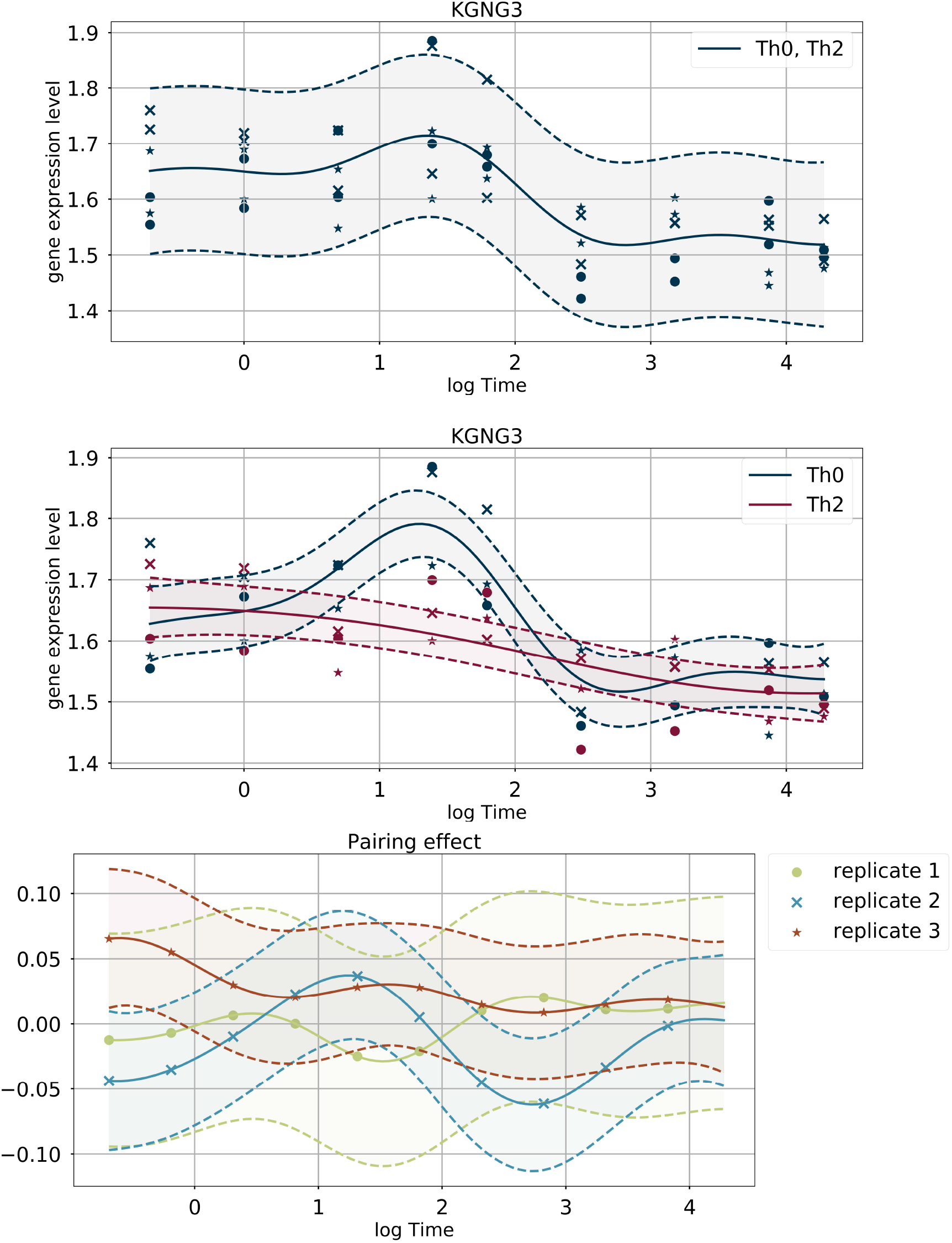
(top) The result of the base model on the gene *KCNG3* (probe set *l552897_a_at*). (middle) The result of the pairing effect model on the gene *KCNG3* and (bottom) the relative pairing effect learned from the data.

**Figure 4:**
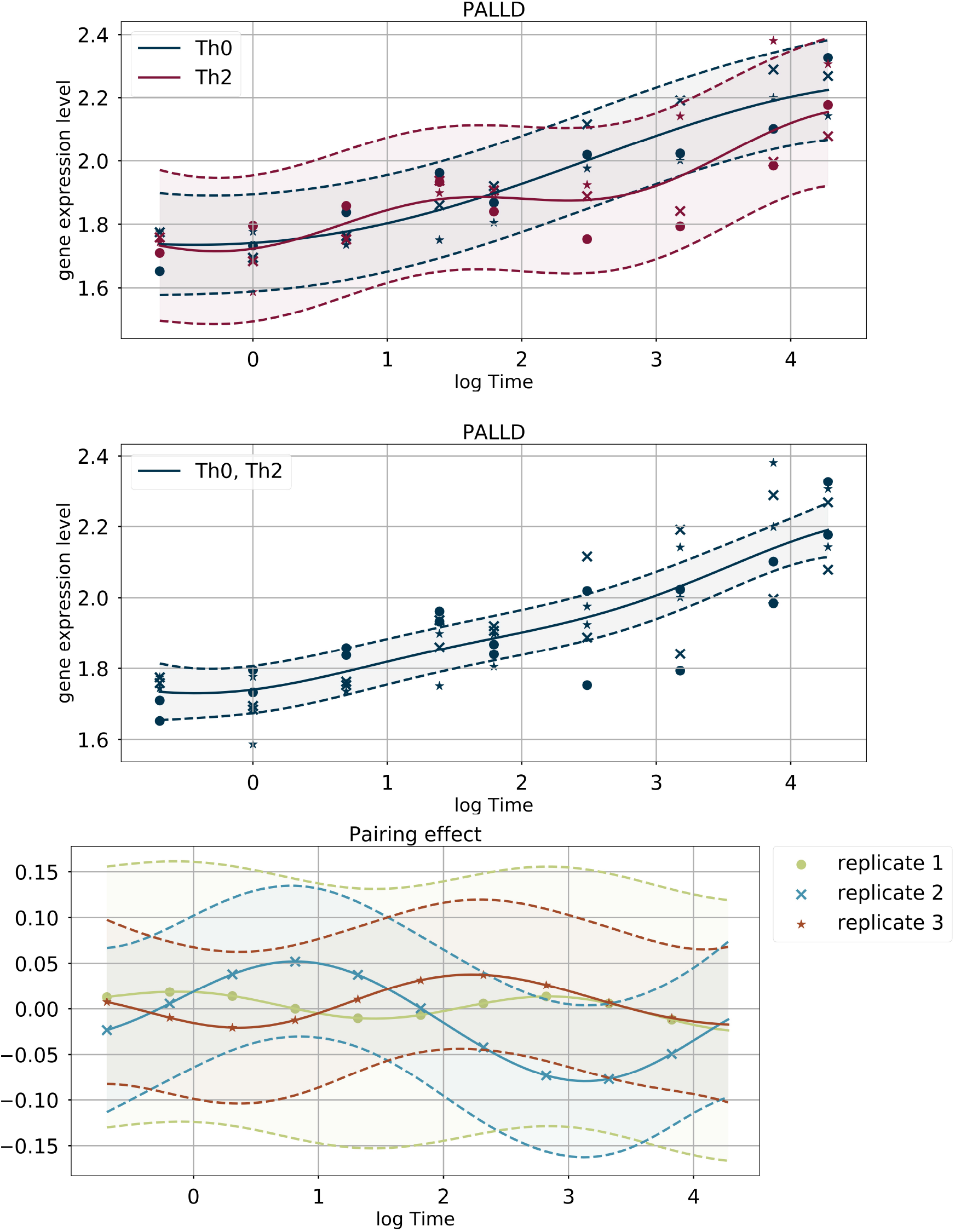
(top) The result of the base model on the gene *PALLD* (probe set *200897_s_at*). (middle) The result of the pairing effect model on the gene *PALLD* and (bottom) the relative pairing effect learned from the data.

#### 3.3 RNA-seq data

**Table 2:** Most frequent partitions obtained by fitting the base model and the pairing effect model on all the genes from the T-helper cell RNA-seq data set. The percentage of the total amount of genes is reported.

| Partition | Base | Pairing |
| --- | --- | --- |
| $\{\{\text{Th0}\}, \{\text{Th17}, \text{Th17+IL1b}, \text{Th17+IL21}, \text{Th17+IL1b+IL21}\}\}$ | 47.9 | 22.8 |
| $\{\{\text{Th0}, \text{Th17}, \text{Th17+IL1b}, \text{Th17+IL21}, \text{Th17+IL1b+IL21}\}\}$ | 9.6 | 19.4 |
| $\{\{\text{Th17}\}, \{\text{Th0}, \text{Th17+IL1b}, \text{Th17+IL21}, \text{Th17+IL1b+IL21}\}\}$ | 0.5 | 6.6 |
| $\{\{\text{Th0}\}, \{\text{Th17}\}, \{\text{Th17+IL1b}, \text{Th17+IL21}, \text{Th17+IL1b+IL21}\}\}$ | 19.7 | 5.6 |
| $\{\{\text{Th0}, \text{Th17}\}, \{\text{Th17+IL1b}, \text{Th17+IL21}, \text{Th17+IL1b+IL21}\}\}$ | 17.4 | 3.9 |
| $\{\{\text{Th0}\}, \{\text{Th17}, \text{Th17+IL21}\}, \{\text{Th17+IL1b}, \text{Th17+IL1b+IL21}\}\}$ | 2.3 | 3.8 |
| others | 2.6 | 37.9 |

**Figure 5:**
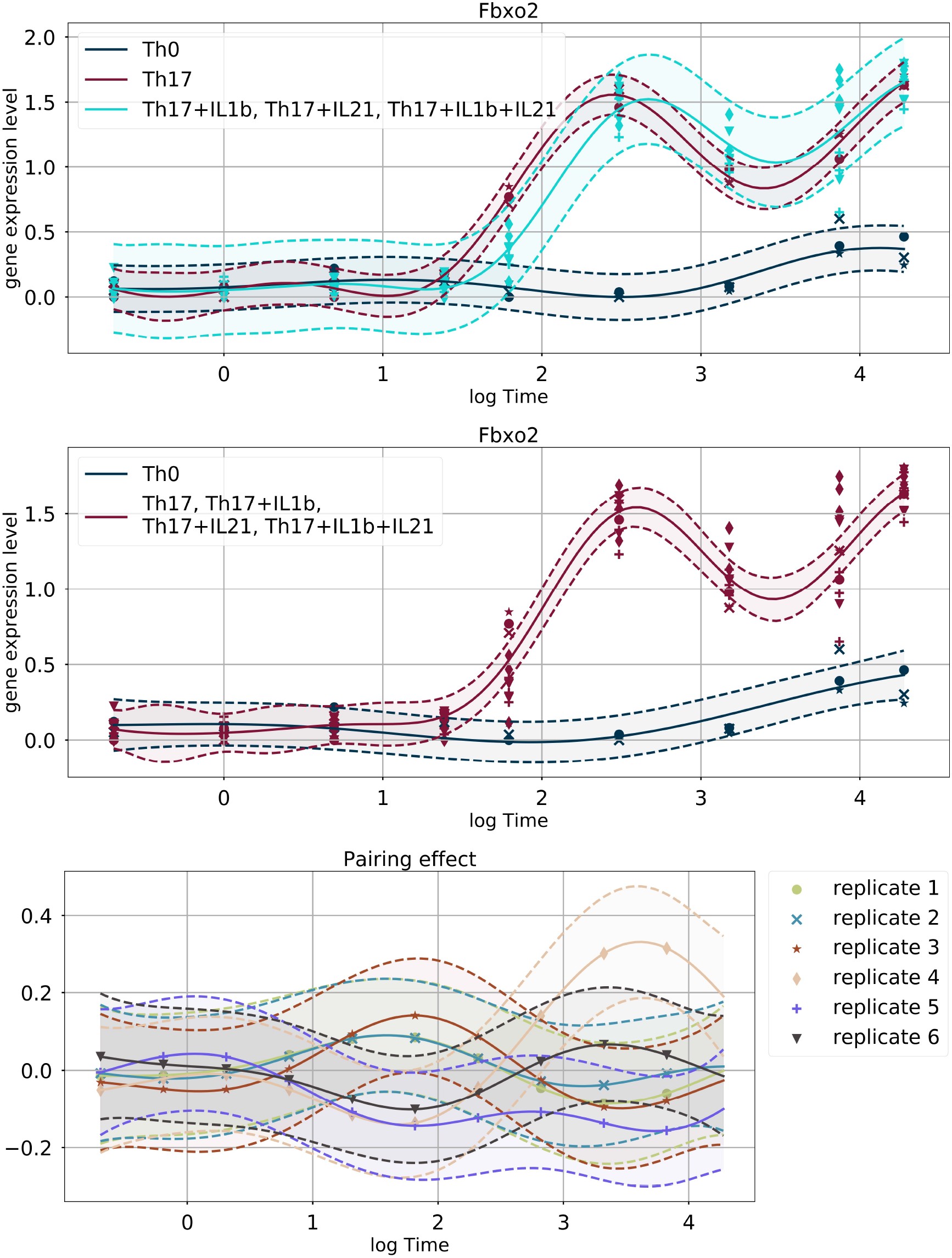
(top) The result of the base model on the gene *Fbxo2*. (middle) The result of the pairing effect model on the gene *Fbxo2* and (bottom) the relative pairing effect learnt from the data.

**Figure 6:**
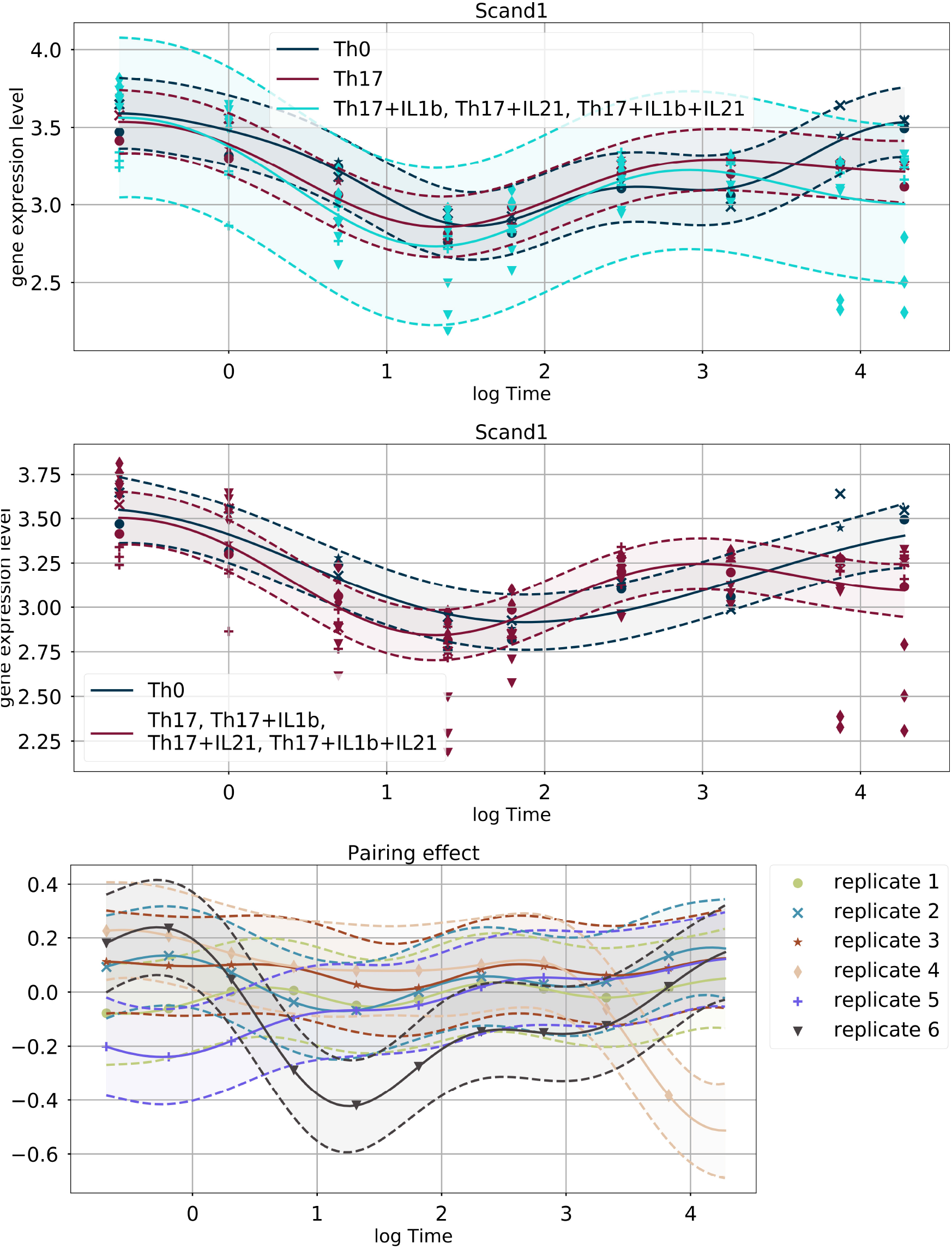
(top) The result of the base model on the gene *Scand1*. (middle) The result of the pairing effect model on the gene *Scand1* and (bottom) the relative pairing effect learnt from the data.

